# Single-cell multiomic QTL mapping reveals state-dependent genetic regulation and associated gene during cellular senescence

**DOI:** 10.64898/2026.09.17.752532

**Authors:** Xianfu Yi, Xueqi Wang, Kunlun Liu, Lin Zhao, Fa Chen, Qianting Jian, Jianhua Wang, Hao Lu, Wei Dong, Yao Zhou, Ying Chang, Xiaoqiong Gu, Pak Chung Sham, Dandan Huang, Mulin Jun Li

## Abstract

How cellular senescence reshapes inherited regulatory effects across chromatin and transcription, and eventually contribute to non-coding risk loci of complex diseases, remains elusive. We generated single-cell multiome ATAC-RNA profiles from proliferating and replicatively senescent primary HUVECs derived from 100 genotyped donors. Alongside state-resolved eQTL and caQTL mapping, we implemented covariance-aware multivariate QTL mapping to identify chromatin accessibility-expression QTLs (caeQTLs), modeling accessibility and expression as a joint phenotype. Joint analysis recovered moderate, asymmetric and modality-distributed associations missed by single-modality scans, revealed senescence-associated regulatory programs, and prioritized candidate causal variants and effector genes at cardiovascular loci. Fine-mapping, chromatin-state annotation and heritability enrichment, together with independent SNP-to-gene algorithms, provided convergent orthogonal support. Mechanistic dissection revealed that rs2019090 modulates *PDGFD* expression and endothelial phenotypes through senescence-amplified, allele-specific, HMGA1-associated enhancer-promoter regulation. This study establishes senescence-resolved joint multiomic QTL mapping as a framework for narrowing the missing regulation gap in complex disease genetics.

## Introduction

Genome-wide association studies have identified thousands of loci for complex traits and diseases, yet most associated variants reside outside protein-coding sequences and do not directly reveal their effector genes or relevant cellular contexts^1,2^. *Cis*-regulatory element (CRE) to gene maps and integrative variant-to-gene linking strategies have substantially improved disease/trait-causal gene interpretation, but their incomplete recall and limited concordance leave a persistent regulatory interpretation gap^3–6^. Molecular QTL mapping provides an orthogonal route to connect inherited variation with intermediate molecular phenotypes. Population-scale studies have increasingly profiled gene expression, chromatin accessibility, histone modifications and chromatin interactions across matched donor cohorts, typically mapping eQTLs, caQTLs and other molecular QTLs separately before integrating them through colocalization, mediation or causal-direction analyses^7–11^. Moreover, conventional eQTLs favor proximal, simple-regulation genes, missing distal variants within complex *cis*-regulatory landscapes^12,13^. Chromatin and histone QTLs capture these upstream effects, often buffered or context-dependent, boosting GWAS colocalization over 2-fold and annotating ∼50% of disease/trait-associated loci^14,15^. However, separate mapping of molecular QTLs followed by *post hoc* integration ignores cross-modal residual covariance. It also requires effects in each modality to pass independent significance thresholds, limiting sensitivity to genetic effects distributed across accessibility and expression^10,16,17^. Whether these paired molecular phenotypes can instead be modeled jointly to improve regulatory QTL discovery remains insufficiently explored.

The regulatory effects of genetic variants are strongly dependent on cellular context. Population-scale single-cell studies have shown that molecular QTL effects can be restricted to particular cell types, activation states and differentiation trajectories, while continuous cell-state modeling can reveal disease-relevant associations that are obscured in bulk tissues or discrete cellular classifications^18–23^. Context-specific QTL studies across endodermal, cardiac and heterogeneous iPSC differentiation have identified genetic effects that emerge, disappear or change magnitude along developmental trajectories^24–26^. Complementary studies of pathogen stimulation and environmental exposures have uncovered response QTLs absent at homeostasis, with continuous modeling of heterogeneous perturbation states further increasing their detection^8,27,28^. Emerging same-cell multiomic studies have further connected context-dependent caQTL and eQTL effects to disease loci in various human tissues^29–31^, supporting the value of tracing genetic effects across coupled regulatory layers. Together, existing studies indicate that context-resolved molecular QTLs can expose otherwise latent regulatory effects and clarify how disease variants propagate through cellular programs. These findings raise the question of whether cellular senescence similarly reshapes the regulatory effects of inherited variation. Although aging-focused multiomic atlases have characterized age-associated changes in gene expression, chromatin accessibility and regulatory coupling^32,33^, whether senescence unmasks, amplifies or attenuates inherited effects across these coupled molecular layers remains unclear.

Here, we generated a donor-scale, state-resolved single-cell multiome resource from proliferating and replicatively senescent human umbilical vein endothelial cells (HUVECs) derived from 100 genotyped donors. Beyond conventional eQTL and caQTL mapping, we established covariance-aware multivariate QTL mapping, treating chromatin accessibility and gene expression as a bivariate molecular phenotype rather than independent traits. By testing genotype-associated displacement of this joint phenotype while accounting for cross-modal residual covariance, this framework captures moderate, asymmetric or modality-distributed effects that may remain subthreshold in single-modality scans. Integrating joint QTLs with cross-state genetic analyses, fine-mapping, regulatory annotations and cardiovascular GWAS, we identify senescence-associated chromatin-expression associations that expand the molecular annotation of cardiovascular disease (CVD) loci and reveal regulatory programs missed by conventional QTL maps. Functional dissection of the rs2019090-HMGA1-*PDGFD* axis further demonstrates how senescence amplifies an inherited, allele-specific regulatory effect from enhancer activity to target-gene expression and endothelial phenotype. Together, this study establishes a framework for tracing context-dependent genetic regulation across molecular layers and narrowing the missing-regulation gap in cardiovascular GWAS.

## Results

### State-resolved conventional eQTL and caQTL mapping establishes a baseline landscape during endothelial senescence

To establish an interpretable single-modality reference for subsequent cross-modal analyses, we first treated the RNA and ATAC measurements generated by the 10x Genomics Multiome assay as separate donor-level molecular phenotypes. Primary HUVECs from 100 genotyped donors were profiled in proliferating and replicatively senescent states following serial passaging (Fig. 1A; see details in Methods). This system recapitulated the progressive acquisition of endothelial replicative senescence and provided a physiologically relevant, cell-type-resolved model for investigating its genetic regulation. Profiling a donor cohort minimized tissue-based sampling biases arising from variable endothelial abundance, incomplete cell recovery and inter-individual differences in cell yield, while enabling controlled and cost-effective donor-level QTL mapping. After modality-specific quality control, 42,671 RNA cells and 42,317 ATAC cells were retained across two independent multiome replicates for each state. Dimensionality reduction revealed broadly homogeneous endothelial populations, with broader dispersion among senescent cells, particularly in the ATAC representation (Fig. S1A). Genotype-based demultiplexing with Vireo^34^ assigned 83.7% and 74.03% of cells to individual donors in the two states, a coverage comparable to or exceeding that in many population-scale single-cell QTL studies (Fig. 1B; see details in Methods). To preserve the donor as the independent unit of genetic association, cells were aggregated within each donor and state before conventional *cis*-eQTL and *cis*-caQTL mapping (Fig. 1C and Fig. S1B; Supplementary Tables S1-S4; see details in Methods). In proliferating and senescent cells, respectively, we identified 647 and 471 eGenes and numerous significant variants and variant-eGene associations. The corresponding caQTL analyses identified 412 and 1,453 caPeaks and numerous variants and variant-caPeak associations (Fig. S1C). Because the two states were profiled independently and differed in cell yield, peak space and testing burden, QTL counts were interpreted descriptively rather than as evidence of differential genetic regulation.

**Fig. 1.**
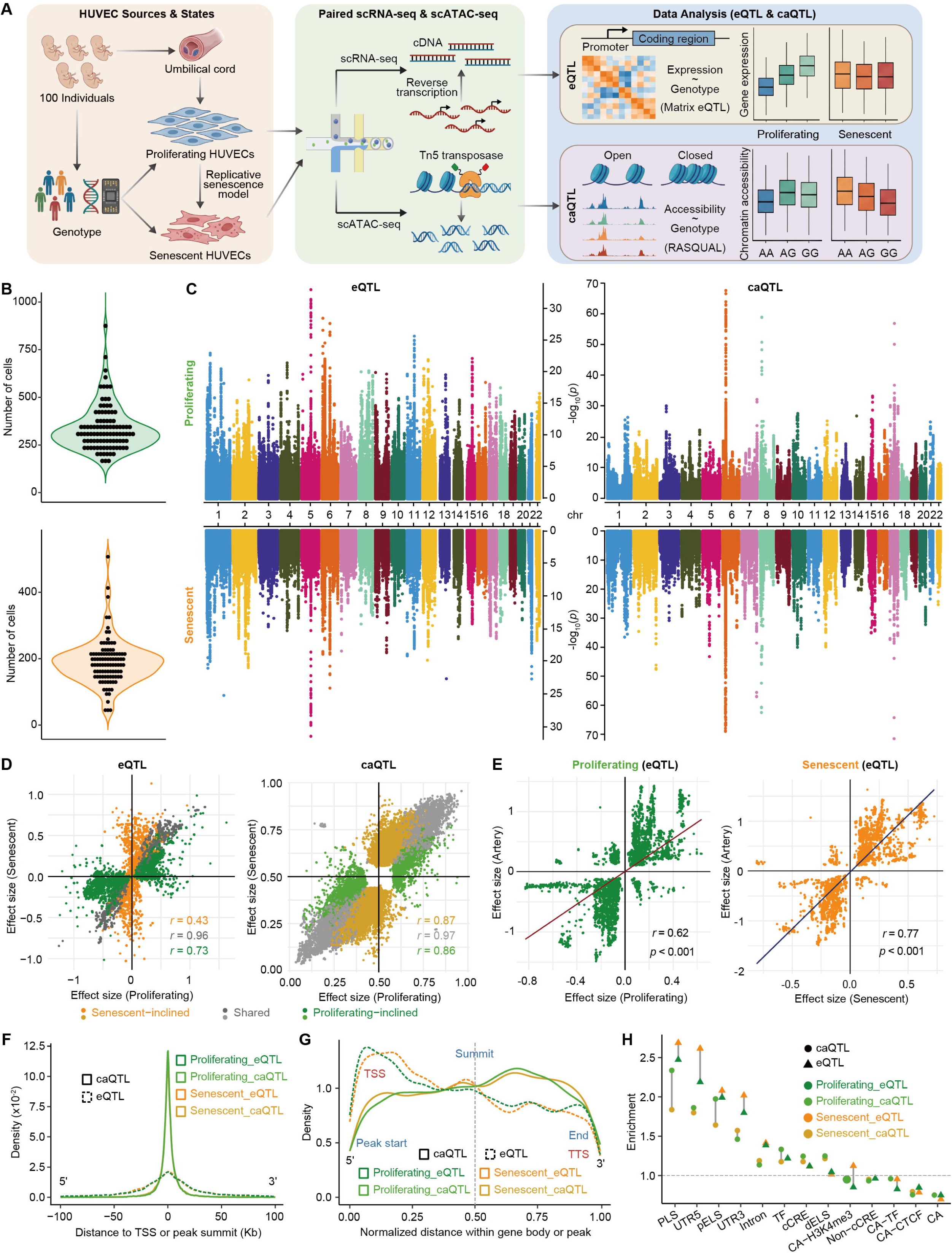
Single-cell eQTL and caQTL mapping establishes a baseline landscape during endothelial senescence. **A**, Experimental design. Primary HUVECs isolated from 100 genotyped donors were subjected to serial passaging to derive proliferating and replicatively senescent states, profiled by paired single-cell RNA-seq and ATAC-seq (10x Genomics Multiome), and analysed for *cis*-eQTLs and *cis*-caQTLs after donor demultiplexing and pseudo-bulk aggregation. **B**, Number of cells confidently assigned to each donor by Vireo demultiplexing in proliferating and senescent states. **C**, Genome-wide Manhattan plots of significant eQTLs (left) and caQTLs (right) in proliferating (top) and senescent (bottom) cells. **D**, Concordance of effect sizes (beta) between proliferating and senescent states for eQTLs (left) and caQTLs (right). **E**, Comparison of HUVEC eQTL effect sizes with GTEx Artery-Aorta eQTLs in proliferating (left) and senescent (right) cells. **F**, Enrichment of eQTL and caQTL variants around transcription start sites (TSS) and ATAC peak summits. **G**, Relative positions of eQTL and caQTL variants within gene bodies and accessible peaks. **H**, Enrichment of eQTL and caQTL variants across *cis*-regulatory element (ENCODE V4 cCRE) and gene structure classes. cCREs were classified as promoter-like signatures (PLS), proximal enhancer-like signatures (pELS), distal enhancer-like signatures (dELS), CTCF-bound (CA-CTCF), H3K4me3-modified (CA-H3K4me3), TF-bound (CA-TF), chromatin-accessible-only (CA) or TF-only regions, or as non-cCRE.

Internal and external benchmarks supported the robustness and biological coherence of these maps. QTLs shared between states showed highly concordant effects, with correlations of 0.96 for eQTLs and 0.97 for caQTLs (Fig. 1D). In GTEx arterial tissues, 27% and 33% of proliferating and senescent variant-eGene associations were recapitulated, respectively, with concordant effect estimates (r = 0.62 and r = 0.77; Pearson’s correlation test); similar patterns were observed in coronary and tibial arteries (Fig. 1E and Fig. S1D,E). Conversely, only a small proportion of GTEx arterial eQTLs were detected in HUVECs, consistent with cell-type and state specificity as well as differences in statistical power (Fig. S1F,G). Consistent with previous findings^14,15^, functional annotation further showed that eQTL variants were enriched near transcription start sites (TSSs) and in promoter-proximal elements, whereas caQTL variants localized preferentially to ATAC peak summits and distal enhancer-like elements (Fig. 1F-H). Despite this validation, overlap between states was limited to 162 eGenes and 304 caPeaks, with low Jaccard coefficients across feature, variant and association levels (Fig. S1C,H). As significance-based overlap is sensitive to differential power, shared and state-inclined QTLs were treated as operational categories. Cochran’s Q heterogeneity tests provided partial support for these assignments, with concordance varying by effect magnitude (Fig. S2A-F).

### Dissecting patterns of genetic regulation across cellular contexts and molecular modalities

To characterize the context dependence of genetic regulation, we first compared eQTL and caQTL signals between proliferating and senescent HUVECs using cross-state colocalization. Among the eQTLs, 130 loci met the predefined criteria for shared genetic signals between states (PP4 ≥ 0.75 and PP4/PP3 ≥ 3), of which 121 (93.08%) corresponded to eQTLs detected in both states (Fig. 2A; Supplementary Table S5). For example, the *LAMC2* locus showed highly concordant effects at rs2276543 in proliferating and senescent cells, whereas most state-inclined eQTLs, including the senescence-associated *PDGFD* rs2019090 locus, did not show evidence for a shared causal signal across states. A similar pattern was observed for caQTLs, with 257 cross-state colocalized loci, 87.94% of which were detected in both states (Fig. 2B; Supplementary Table S6). Although shared QTLs were more likely to colocalize across states, approximately one-quarter of shared eQTLs and caQTLs did not meet the colocalization criteria, consistent with allelic heterogeneity, linkage disequilibrium structure or limited statistical power. Conversely, only 1.32% of state-inclined eQTLs and 5.54% of state-inclined caQTLs showed significant cross-state colocalization. Representative loci illustrated both highly concordant signals and state-preferential associations across the two molecular modalities (Fig. 2A,B and Fig. S3A,B). Thus, cross-state colocalization provided locus-level support for the genetic continuity of most shared QTLs, while prioritizing a distinct subset of associations whose molecular effects were preferentially manifested in proliferating or senescent cells.

**Fig. 2.**
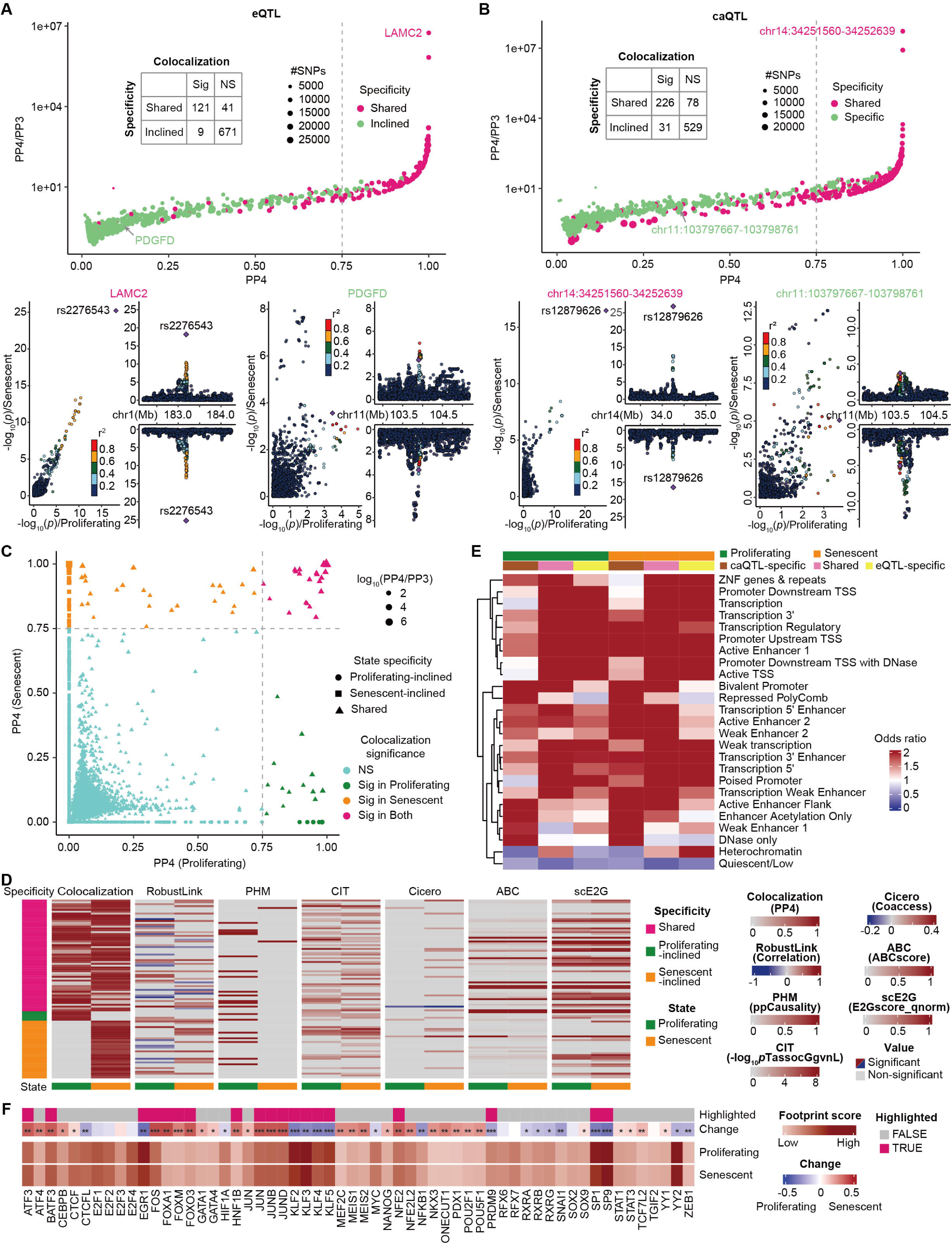
Comparative analysis of eQTLs and caQTLs across states and molecular modalities. **A**, Cross-state colocalization of eQTLs. The scatter plot summarizes colocalization across all eQTLs; below, a state-shared and colocalized example (*LAMC2*, rs2276543) and a senescence-inclined example (*PDGFD*, rs2019090). **B**, As in **A**, for caQTLs. Representative peaks are chr14:34251560-34252639 (rs12879626; shared and colocalized) and chr11:103797667-103798761 (rs2019090; senescence-inclined). **C**, Summary of eQTL-caQTL colocalization across states, showing the proportions of relationships colocalized in both states, one state or neither. **D**, Multi-dimensional evidence support (RobustLink, PHM, CIT, Cicero, ABC, scE2G) for colocalized peak-gene pairs, and agreement between colocalization class and evidence class. **E**, Enrichment of eQTL and caQTL variants across the 25 Roadmap HUVEC (E122) chromatin states. **F**, Differential transcription-factor binding between proliferating and senescent cells from JASPAR 2024 motifs (TOBIAS footprinting). Significance (\*\*\**p* <1e-200, \*\**p* <1e-150, \**p* <1e-100) and the 95th/5th-percentile change thresholds are defined in Methods.

We next examined whether eQTLs and caQTLs capture distinct layers of genetic regulation. Across both states, caQTL variants were located farther from TSSs than eQTL variants and showed limited overlap with eQTL variants, with low Jaccard coefficients in proliferating and senescent cells (Fig. S3C,D). Variants detected by both modalities showed the highest proportion of causal-inference support via CIT^35^, whereas caQTL-specific variants showed slightly greater causal-inference support than eQTL-specific variants, suggesting that accessibility QTLs capture regulatory signals not readily detected at the transcriptional level. Direct eQTL-caQTL colocalization was nevertheless uncommon in our data, with only 96 of 59,518 tested QTL relationships reaching the predefined threshold in at least one state (Fig. 2C; Supplementary Tables S7,S8). Among these, 21 relationships colocalized in both states, whereas additional relationships were specific to proliferating or senescent cells. 71 of the colocalized peak-gene relationships had support from at least one independent regulatory-linking framework, including RobustLink^36^, PHM^37^, CIT^35^, Cicero^38^, ABC^3^ or scE2G^39^ (Fig. 2D). However, agreement between colocalization and peak-gene annotations was incomplete, particularly for state-shared links, emphasizing that genetic colocalization alone is insufficient to resolve regulatory targets. Consistent with their distinct molecular roles, eQTLs preferentially enriched in transcription-associated chromatin states, whereas caQTLs were enriched in DNase-, enhancer- and Polycomb-associated states (Fig. 2E).

To prioritize putative causal variants and define their regulatory contexts, we fine-mapped caQTLs and eQTLs using CAVIAR^40^, respectively (Supplementary Tables S9 and S10). We identified 369 and 1,718 variant-caPeak associations in proliferating and senescent cells, respectively, with comparable numbers of unique variants showing posterior causal probabilities exceeding 0.8. Correspondingly, 725 and 923 variant-eGene associations yielded high-probability variants. Most features were resolved to no more than two prioritized variants, comprising the large majority of caPeaks and eGenes (Fig. S4A). Among fine-mapped molecular QTL variants, 6.0% in proliferating cells and 6.4% in senescent cells had CIT support, respectively (Fig. S4B). Fine-mapped eQTLs remained enriched in transcription-associated chromatin states, whereas caQTLs preferentially occupied accessible and enhancer-associated states, recapitulating the modality-specific architecture of the full QTL maps (Fig. S4C). Transcription factor (TF) footprinting analysis further revealed state-associated regulatory programs. Analyses using JASPAR 2024^41^ identified preferential activity of KLF4 and SP1 in proliferating cells and FOS and JUN in senescent cells, whereas CTCF, CTCFL and E2F-family factors showed limited state preference (Fig. 2F and Fig. S5A). These patterns were reproduced using HOCOMOCO motifs^42^ (Fig. S5B,C). Together, these analyses prioritized putative causal variants and delineated modality- and state-associated regulatory architectures during endothelial senescence.

### Covariance-aware joint QTL mapping reveals state-resolved chromatin-expression regulatory associations

Separate eQTL and caQTL mapping of single-cell multiome data provided complementary yet fragmented views of genetic regulation. Their distinct genomic distributions and limited cross-modal colocalization arise largely from heterogeneous effect sizes and detection power across molecular layers, leaving weak or asymmetric cross-modal effects systematically undetected. In particular, variants with moderate or asymmetric effects on accessibility and expression may remain undetected when each phenotype is tested separately, whereas *post hoc* integration retains only signals that satisfy modality-specific thresholds^16,17^. Moreover, existing proximity-, correlation- and contact-based CRE-gene maps are not inherently genetically anchored by construction, while colocalization depends on sufficiently strong univariate signals. We therefore modeled the paired donor-level accessibility and expression measurements for each candidate variant-peak-gene triplet as a bivariate phenotype using multivariate multiple regression (MMR). The genotype term was evaluated with Pillai’s trace as joint statistics, accounting for cross-modal residual covariance in a single omnibus test (Fig. 3A and Fig. S6A,B; Supplementary Tables S11 and S12; see details in Methods). We termed significant associations (adjusted using the Benjamini-Hochberg procedure) chromatin accessibility-expression QTLs, or caeQTLs. These associations genetically anchor candidate CRE-gene relationships and capture effects manifested in either or both modalities (caeQTLs denote genetic association, not mediation of expression by accessibility).

**Fig. 3.**
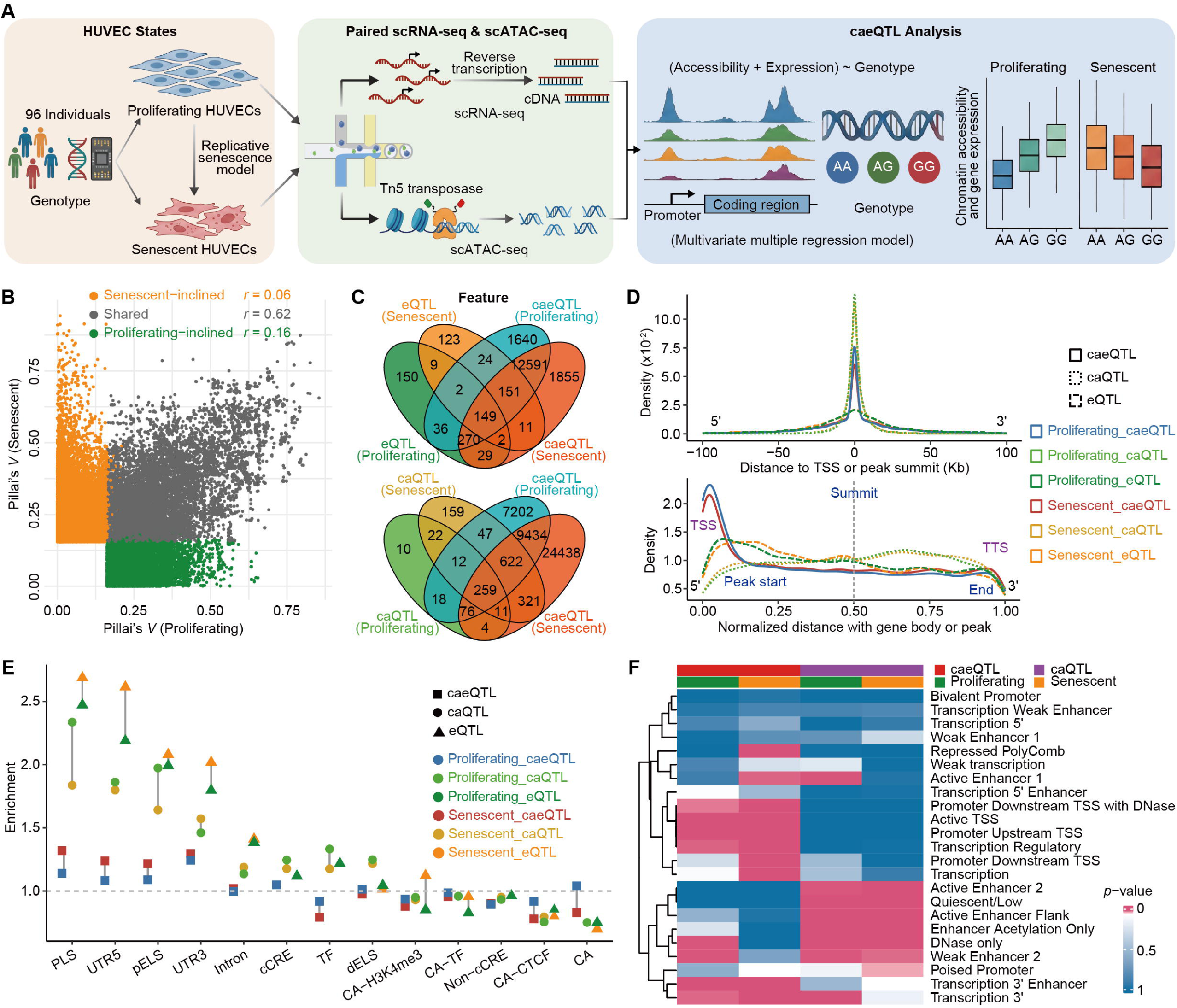
Covariance-aware joint QTL mapping identifies state-resolved chromatin accessibility-expression (caeQTL) associations. **A**, Analytic framework. Matched per-donor gene expression and chromatin accessibility are modelled as a bivariate phenotype by multivariate multiple regression (MMR); the genotype term is tested with Pillai’s trace, accounting for cross-modal residual covariance. **B**, Concordance of the joint statistic (Pillai’s trace *V*) between states for caeQTLs; Pillai’s trace *V* (0 ≤ V ≤ 1) is a non-directional joint effect statistic, not a signed regression coefficient. **C**, Feature-level overlap of caeQTLs with eQTLs (top; eGene level) and caQTLs (bottom; caPeak level). **D**, Enrichment of caeQTL, eQTL, and caQTL variants around TSS and peak summits (top) and their relative positions within gene bodies and accessible peaks (bottom). **E**, Enrichment of caeQTL, eQTL, and caQTL variants across cis-regulatory element (ENCODE V4 cCRE) and gene structure classes. cCREs were classified as promoter-like signatures (PLS), proximal enhancer-like signatures (pELS), distal enhancer-like signatures (dELS), CTCF-bound (CA-CTCF), H3K4me3-modified (CA-H3K4me3), TF-bound (CA-TF), chromatin-accessible-only (CA) or TF-only regions, or as non-cCRE. **F**, Enrichment of caeQTL and caQTL variants across the 25 Roadmap HUVEC (E122) chromatin states.

We identified 132,393 and 231,506 significant caeQTL associations in proliferating and senescent cells, respectively, encompassing large numbers of variants and peak-gene features (Fig. S6C). Leveraging covariance, the joint test mostly detected associations significant in only one or neither modality alone, but cross-state overlap was limited (Fig. S6D). Nevertheless, shared associations showed substantially greater cross-state concordance in Pillai’s trace statistic (*V*) than state-inclined associations (r = 0.62 versus r = 0.06 and .r= 0.16; Fig. 3B). At the feature level, joint mapping recovered 70.63% and 66.45% of eGenes and 88.59% and 83.48% of caPeaks identified in proliferating and senescent cells, respectively, whereas overlap at the exact variant and association levels was still considerably lower (Fig. 3C and Fig. S7A-C). Conversely, when projected onto the corresponding single-modality spaces, caeQTLs captured a large fraction of both caQTLs and eQTLs. Most joint associations were not individually significant in either univariate scan, and those with univariate support were predominantly accessibility associated (Fig. S8A-C), consistent with the recovery of asymmetric or distributed cross-modal effects. The excess of small joint *p*-values and preferential detection of associations carrying moderate component signals further supported increased sensitivity beyond *post hoc* intersection (Fig. S8D,E).

The caeQTL map exhibited a coherent regulatory architecture distinct from that of the single-modality maps. Compared with eQTL and caQTL variants, caeQTL variants were more concentrated immediately upstream and downstream of TSSs and less represented within gene bodies, with a less distal distribution than caQTL variants (Fig. 3D and Fig. S9A). They were preferentially enriched in promoter-like sequences, 5′ untranslated regions and proximal enhancer-like elements, with stronger enrichment of several transcriptionally engaged annotations in senescent cells (Fig. 3E). Roadmap chromatin-state analyses similarly placed caeQTLs in active TSS, transcriptional and enhancer-associated states, whereas caQTLs occupied predominantly accessibility-associated states (Fig. 3F). Genes preferentially highlighted by joint rather than expression-only evidence were enriched in immune and adhesion pathways, whereas the broader caeQTL-linked gene set captured cellular senescence, longevity-regulating and cell-cycle programs that were weakly represented among conventional eQTL genes (Fig. S9B,C). Approximately 40% of linked genes were also represented among arterial GTEx eGenes, although resolution differed at the variant-gene level (Fig. S9D,E). Finally, scE2G support and gene expression were prominent predictors of caeQTL status, whereas nucleosome depletion and chromatin accessibility showed state-dependent contributions (Fig. S9F). Thus, joint mapping preferentially identified genetically associated, transcriptionally engaged regulatory elements linked to senescence-relevant cellular programs.

We then evaluated caeQTLs using convergent genetic, epigenomic and regulatory evidence. Across nine orthogonal evidence sources (ABC, Cicero, CIT, colocalization, PHM, RobustLink, scE2G, Hi-C and Micro-C), the evidence-supported caeQTL subset was characterized by greater overlap with causal SNP-to-gene annotations than unsupported associations (Fig. 4A,B; Supplementary Table S13). After conditioning on both caQTL and eQTL annotations, senescent-state evidence-supported caeQTLs remained significantly enriched for CAD heritability (Fig. 4C,D), indicating regulatory information not subsumed by either univariate map; unconditioned enrichment was comparable between groups. Joint mapping recovered 77.97% and 64.55% of eQTL-caQTL colocalized relationships in proliferating and senescent cells, respectively (Fig. 4E). It additionally identified 104,432 associations not detected by colocalization. These caeQTL-specific associations retained ABC and RobustLink support comparable to colocalized associations (Fig. 4F and Fig. S10A), indicating that the added sensitivity reflects association-level power rather than weaker evidence. They also showed more frequent additional support than associations prioritized by CIT or PHM alone, including a higher proportion supported by multiple evidence classes (Fig. 4G). CAVIAR-based locus prioritization further nominated 5,562 candidate variants, of which 64.3% belonged to significant caeQTLs and 10.9% had CIT support (Fig. S10B; Supplementary Tables S14 and S15). Collectively, these analyses establish caeQTL mapping as a systematic approach for expanding genetically anchored CRE-gene prioritization, particularly for modest and senescence-associated effects.

**Fig. 4.**
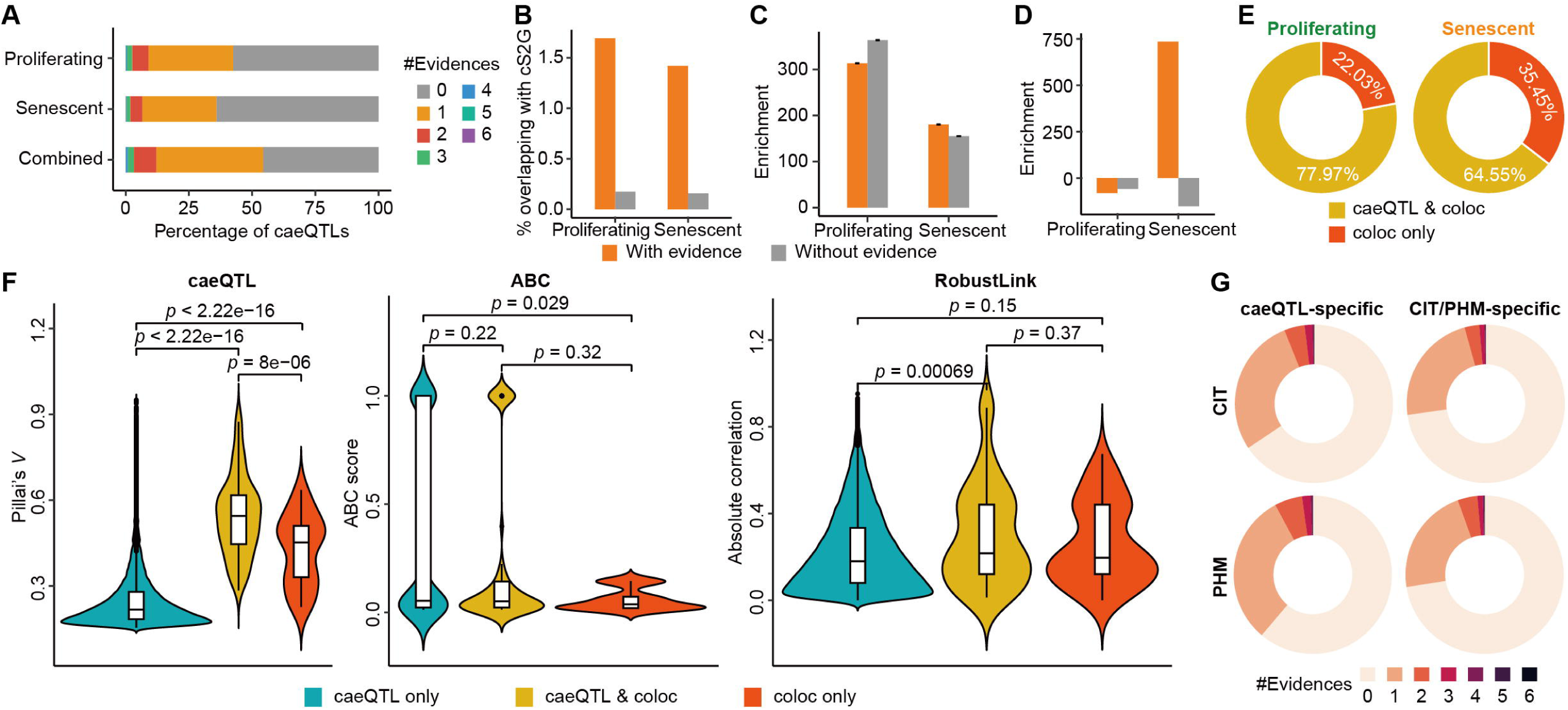
Multi-dimensional evidence support for caeQTLs. **A**, External evidence support across nine computational and experimental sources (ABC, Cicero, CIT, colocalization, Hi-C, Micro-C, PHM, RobustLink, and scE2G). **B**, Overlap of evidence-supported versus unsupported caeQTLs with causal SNP-to-gene pairs (cS2G). **C**, S-LDSC heritability enrichment of evidence-supported versus unsupported caeQTLs (unconditioned). **D**, As in **C**, after conditioning on caQTL and eQTL annotations; senescent-state evidence-supported caeQTLs retain significant enrichment, indicating information not subsumed by either univariate map. **E**, Correspondence between significant eQTL-caQTL colocalization and caeQTLs in proliferating (left) and senescent (right) cells. **F**, Comparison of caeQTL-specific, shared, and colocalization-specific associations in joint effect magnitude (left), ABC score (middle), and RobustLink correlation (right). **G**, Proportion of caeQTL-specific associations with multi-dimensional evidence relative to those prioritized by CIT or PHM alone.

### Joint QTL mapping prioritizes disease-relevant regulatory variants and genes

We next asked whether joint chromatin-expression QTLs provide complementary information for interpreting disease-associated genetic variation. Across GWAS loci representing eight disease systems from our CAUSALdb^2^, CVD showed the strongest overlap with all three QTL classes, but the overlap was consistently higher for caeQTLs than for conventional caQTLs or eQTLs in both proliferating and senescent cells (Fig. 5A and Fig. S11A). Across the eight systems, at least one QTL class covered 8.36% of GWAS variants in proliferating cells and 14.5% in senescent cells, with the higher coverage in senescence consistent with the broader state-resolved regulatory landscape (Fig. S11B). Most GWAS-overlapping variants were assigned to only one QTL class, with 88.0% and 91.4% belonging exclusively to caQTLs, eQTLs or caeQTLs in the two states, respectively, indicating that the three molecular phenotypes capture largely non-redundant genetic signals for complex diseases. This complementarity was particularly evident for cardiovascular traits. The fraction of CVD-associated variants unique to caeQTLs reached 18.4% in proliferating cells and 16.61% in senescent cells, exceeding the corresponding fractions observed for the other disease systems (Fig. 5B). The senescent-state caeQTLs overlapped CVD GWAS variants more than proliferating-state caeQTLs, and overlap was highest for caeQTLs shared between states (Fig. 5C), defining a robust core of condition-independent cardiovascular signals. Among GWAS-overlapping variants, cS2G^4^ support was strongest for variants classified as both caQTL and eQTL in only one state, closely followed by state-resolved caeQTL groups and eQTL groups (Fig. 5D and Fig. S11C). The preferential overlap of senescence-associated caeQTLs with cardiovascular GWAS signals suggests that endothelial senescence can reveal regulatory effects absent from proliferating-cell and steady-state QTL resources, thereby providing complementary molecular annotation for cardiovascular loci^43^ and helping to address the missing regulation gap in GWAS interpretation.

**Fig. 5.**
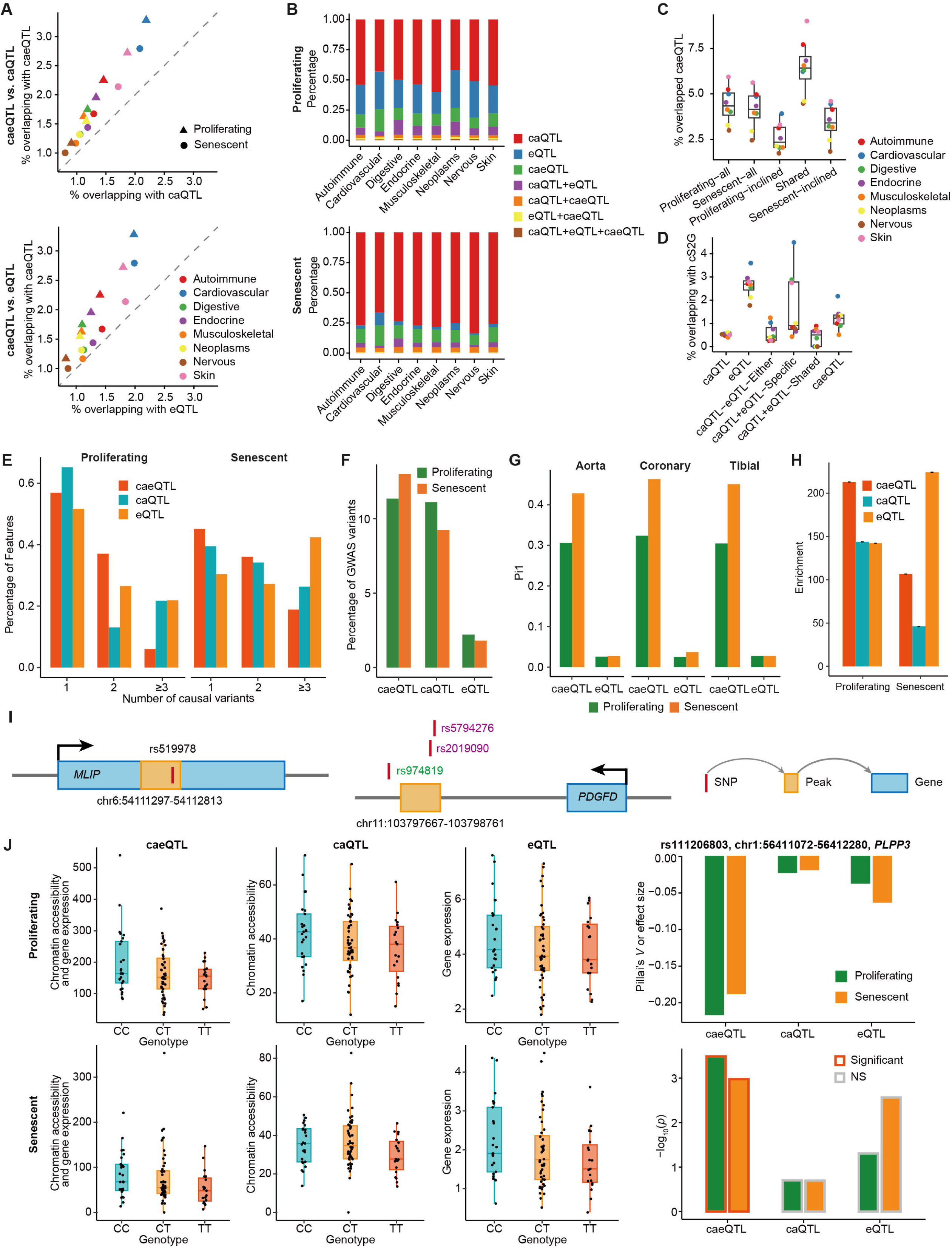
caeQTL-based detection of disease-relevant genetic variants. **A**, Overlap of caeQTLs versus caQTLs (top) and eQTLs (bottom) with GWAS variants across eight disease systems; CVD shows the strongest overlap, consistently higher for caeQTLs. **B**, Composition of GWAS-overlapping QTL variants by caeQTL/caQTL/eQTL class across systems. **C**, Overlap with GWAS variants by state-specificity group of caeQTLs. **D**, cS2G support for GWAS-overlapping QTL variants across QTL classes and state categories. **E**, Number distributions of prioritized causal variants. **F**, Reproduction rate of prioritized variants against CVD GWAS credible sets. **G**, Sharing of HUVEC caeQTLs and eQTLs with GTEx arterial eQTLs (Storey’s π_1_). **H**, S-LDSC heritability enrichment of caeQTL, caQTL, and eQTL variants for CVD. **I**, Representative loci comparing caeQTL, caQTL and eQTL colocalization with CVD GWAS (black, variants shared by caeQTL and colocalization; green, colocalization-only; purple, caeQTL-only). Joint colocalization prioritized *MLIP* in proliferating cells and *PDGFD* in senescent cells. **J**, Effect size and significance of the rs11206803-*PLPP3* link across caeQTL, caQTL, and eQTL analyses; the link is detected by joint mapping in both states but is not significant in either univariate scan. caeQTL *p*-values were derived from the approximate F-test based on Pillai’s trace.

To further evaluate the resolution and disease relevance of the three QTL classes, we compared their CAVIAR-based variant prioritization and relationship to CVD GWAS credible sets. caeQTL associations were generally resolved to a small number of putative causal variants. Prioritization narrowed most caeQTL signals to one or two candidate variants, accounting for 93.98% of associations in proliferating cells and 81.16% in senescent cells, compared with lower proportions for caQTLs and eQTLs (Fig. 5E). caeQTL-prioritized variants also overlapped CVD GWAS credible sets more frequently than eQTL-prioritized variants and at rates comparable to or higher than caQTLs (Fig. 5F). The three QTL classes showed limited overlap at CVD loci, indicating that they capture complementary regulatory signals (Fig. S11D-G). Consistently, HUVEC caeQTLs showed greater sharing with arterial GTEx eQTLs than conventional HUVEC eQTLs, particularly in senescent cells (Fig. 5G). These findings indicate that the disease relevance of genetic regulation differs across molecular modalities and cellular states. caeQTLs showed consistently strong enrichment for CVD-associated variants in both states, whereas caQTLs showed weaker enrichment and eQTLs varied substantially by state (Fig. S11H). Notably, state-inclined caeQTLs were more strongly associated with CVD than shared caeQTLs, highlighting the disease relevance of context-dependent joint regulation (Fig. S11I). S-LDSC analysis further showed greater CVD heritability enrichment for caeQTLs in proliferating cells and for eQTLs in senescent cells (Fig. 5H). Together, these results demonstrate that disease-associated regulatory variation is not uniformly represented across molecular layers or cellular states and that state-resolved joint QTLs prioritize disease-relevant signals that remain inconspicuous in conventional single-modality maps.

We further assessed the capacity of caeQTLs to prioritize regulatory targets supported by convergent disease and molecular genetic evidence. First, joint colocalization with CVD GWAS signals prioritized *MLIP* in proliferating cells and *ATP13A2*, *MFAP2* and *PDGFD* in senescent cells, all of which were also identified by caeQTL mapping (Fig. 5I; Fig. S12A,B; Supplementary Table S16). Second, we evaluated genes previously implicated in CAD through endothelial CRISPR perturbation studies^44^. Joint mapping recovered several experimentally validated candidates, including *PLPP3*, *PECAM1*, *COL4A2*, *BCAR1*, *BMP1* and *PGF* (Fig. 5J and Fig. S13A-D). As a banner example, the rs11206803-*PLPP3* regulatory link was detected by joint mapping in both states yet was non-significant in either univariate scan, directly illustrating the sensitivity gain of joint testing. Its associated accessible region independently gained H3K4me1 in senescent cells, consistent with enhancer activity. Similarly, the rs4968721-*PECAM1* link was detected only by joint mapping, whereas *COL4A2*, *BCAR1*, *BMP1* and *PGF* were preferentially identified in senescent cells. Together, these results show that caeQTL mapping recover disease-relevant regulatory links missed by single-modality analyses and, supported by convergent genetic and functional evidence, prioritizes senescence-associated loci for mechanistic validation.

### rs2019090 modulates allele-specific chromatin openness and *PDGFD* expression in the course of senescence

Among the disease-relevant loci prioritized above, the rs2019090-*PDGFD* locus exhibited concordant but markedly stronger genetic effects in senescent cells across eQTL, caQTL and caeQTL analyses, prompting its mechanistic investigation as a representative senescence-amplified regulatory locus relevant to cardiovascular disease (Figs. 2A,B and 6A,B and S14A,B). rs2019090 lies within an intron of AP002989.1, 109Lkb downstream of *PDGFD*. Previous work in coronary artery smooth muscle cells established rs2019090 and *PDGFD* as the functional variant and effector gene at the 11q22.3 CAD locus, implicated preferential FOXC1/FOXC2 activity at the risk A allele and linked *PDGFD* to smooth-muscle-cell state transitions and vascular inflammation^45^. We therefore asked a distinct question that how endothelial senescence modifies the regulatory penetrance and molecular wiring of this established CAD locus. During senescence, the rs2019090-containing peak gained chromatin accessibility, H3K27ac and H3K4me1, whereas H3K4me3 remained weak, defining a senescence-responsive enhancer-like element (Fig. 6A and Fig. S14C,D). These changes suggested that senescence potentiates a pre-existing regulatory locus rather than creating an effect *de novo*.

**Fig. 6.**
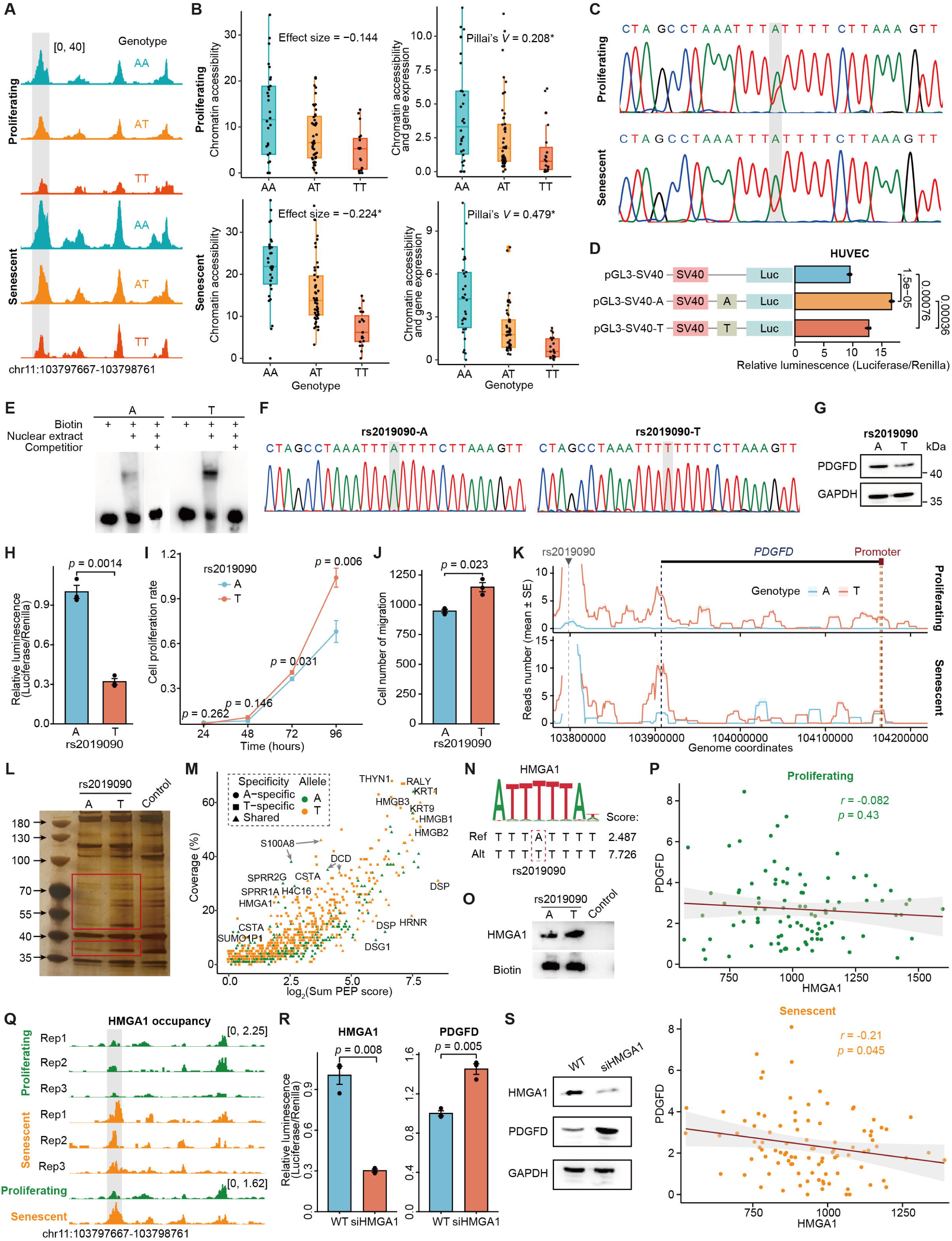
rs2019090 regulates chromatin accessibility and *PDGFD* expression through senescence-amplified, allele-specific enhancer-promoter regulation. **A**, Chromatin accessibility (grey-highlighted peak chr11:103797667-103798761) at the rs2019090 locus across states and genotypes. B, Effect size and significance of rs2019090 in caQTL and caeQTL analyses in proliferating and senescent cells (*, statistically significant). C, Allele-specific chromatin accessibility at rs2019090 by ATAC-PCR in heterozygous cells, confirming greater allelic imbalance in senescent cells. **D**, Dual-luciferase reporter assay showing enhancer activity of the rs2019090 region with stronger activity for the A allele, most pronounced in HUVECs. **E**, EMSA showing stronger total nuclear-protein binding to the T allele. **F**, Single-nucleotide editing of rs2019090 (A to T). **G**, PDGFD expression across genotypes. **H**, RT-qPCR of PDGFD after editing. **I**, Proliferation of HUVECs across genotypes. **J**, Migration of HUVECs across genotypes. K, MNase-based 4C showing a permissive rs2019090-*PDGFD* promoter interaction strengthened in senescence, with a senescence-associated gain of A-allele engagement. L, Allele-specific transcription-factor binding at rs2019090. M, Mass-spectrometry comparison of DNA pull-down products between alleles. **N**, Predicted HMGA1 binding motif flanking rs2019090. O, Allele-specific HMGA1 binding at rs2019090. P, Correlation of *HMGA1* and *PDGFD* expression in proliferating (top) and senescent (bottom) HUVECs; the negative correlation is specific to senescent cells. **Q**, CUT&Tag showing increased HMGA1 occupancy at the rs2019090 enhancer during senescence. R, RT-qPCR of HMGA1 and PDGFD after HMGA1 knockdown. **S**, Western blot of HMGA1 and PDGFD after HMGA1 knockdown.

When effects were oriented to the CAD-risk A allele, rs2019090 was associated with greater chromatin accessibility in both states, whereas its component eQTL and caQTL signals reached FDR significance preferentially in senescent cells (Fig. 6A,B and Fig. S14A,B). The caeQTL remained significant in both states, but the joint effect magnitude (Pillai’s trace statistic *V*) increased from 0.208 in proliferating cells to 0.479 in senescent cells. Allele-specific ATAC-PCR in heterozygous cells independently confirmed greater allelic imbalance in senescent cells (Fig. 6C). Thus, rs2019090 showed a senescence-amplified rather than senescence-specific effect, with allelic differences becoming more pronounced as the enhancer acquired an active chromatin state. Functionally, a dual-luciferase reporter assay demonstrated regulatory activity for a 500-bp fragment surrounding rs2019090, with the A allele driving stronger reporter expression than the T allele. This allele-dependent activity was more pronounced in HUVECs than in 293T cells, supporting both the regulatory function of the locus and its endothelial context dependence (Fig. 6D and Fig. S14E). Electrophoretic mobility shift assay nevertheless revealed stronger total nuclear-protein binding to the T allele (Fig. 6E), suggesting that its lower regulatory activity might reflect preferential recruitment of a repressive factor. Editing the endogenous A allele to T using CRISPR/Cas9 in immortalized HUVECs reduced *PDGFD* expression and increased endothelial proliferation and migration (Fig. 6F-J and Fig. S14F).

To assess direct chromatin contact, we performed MNase-based 4C, which confirmed a permissive interaction between the rs2019090-containing CRE and the *PDGFD* promoter that was strengthened in senescent cells (Fig. 6K and Fig. S15A). Allele-resolved analysis showed that the interaction was predominantly T-derived in proliferating cells, whereas senescence markedly increased the contribution of the A allele, indicating state-dependent changes of enhancer-promoter communication. DNA pull-down coupled with mass spectrometry and allele-specific motif analysis independently converged on HMGA1 as a T-biased binding factor, which was confirmed by immunoblotting (Fig. 6L-O, Fig. S15B,C and Supplementary Tables S17 and S18). HMGA1 is a non-histone AT-hook architectural chromatin factor that can function as a context-dependent transcriptional repressor or buffer through direct promoter inhibition, interference with coactivator recruitment and organization of repressive three-dimensional chromatin environments, including during senescence^46,47^. Interestingly, *HMGA1* and *PDGFD* expression were negatively correlated in arterial GTEx tissues and specifically in senescent HUVECs, but not in proliferating cells (Fig. 6P and Fig. S15D). CUT&Tag further showed increased HMGA1 occupancy at the rs2019090-containing enhancer during senescence, linking its recruitment to the state-dependent regulatory activity of this element (Fig. 6Q). Functionally, HMGA1 depletion increased *PDGFD* transcript and protein abundance, supporting HMGA1 as a negative regulator of the rs2019090-*PDGFD* axis (Fig. 6R,S). Together, these findings support a layered model in which senescence increases accessibility of the rs2019090 enhancer and strengthens its contact with the *PDGFD* promoter, including a marked gain in A-allele engagement. Within this permissive chromatin configuration, preferential HMGA1 binding to the T allele may constrain transcriptional output via promoter contact, whereas the A allele remains comparatively less constrained and supports higher *PDGFD* expression. Thus, senescence amplifies a pre-existing allelic effect by increasing enhancer competence, while allele-selective HMGA1 recruitment shapes transcriptional output.

## Discussion

In this study, we established a donor-scale, state-resolved framework for investigating how cellular senescence reshapes inherited regulatory variation in human endothelial cells. By profiling primary HUVECs from 100 genotyped donors in proliferating and replicatively senescent states using paired single-cell ATAC and RNA sequencing, we generated complementary eQTL and caQTL maps while maintaining a defined cellular context and donor-level genetic resolution. We then implemented covariance-aware multivariate QTL mapping, treating chromatin accessibility and gene expression as a joint molecular phenotype rather than independent traits. This approach expanded the discovery of genetically anchored peak-gene associations, particularly for moderate, asymmetric or modality-distributed effects that were not significant in either single-modality analysis alone. The resulting caeQTL landscape displayed coherent genomic, epigenomic and pathway-level properties, showed enrichment for cardiovascular disease signals beyond conventional QTL annotations, and preferentially highlighted senescence-relevant regulatory programs. Mechanistic dissection of the rs2019090-*PDGFD* locus revealed a senescence-amplified, allele-specific regulatory axis linking enhancer competence and HMGA1 occupancy to enhancer-promoter communication, *PDGFD* expression and endothelial proliferation. These findings establish senescence-resolved joint multiomic QTL mapping as a framework for tracing inherited variation across molecular layers, improving the regulatory interpretation of GWAS signals and prioritizing context-dependent mechanisms for functional investigation.

Using primary HUVECs from 100 donors enabled population-scale analysis of endothelial replicative senescence in a defined cellular context while reducing several sources of heterogeneity inherent to tissue-based single-cell QTL studies. Tissue collection and dissociation can induce cell-type-specific stress programs and alter the recovery and relative representation of endothelial subtypes, thereby affecting the molecular phenotypes and statistical power available for donor-level QTL mapping^48^. A controlled donor-derived system can therefore facilitate the recruitment and profiling of a genetically diverse cohort under a more standardized and potentially cost-efficient design, consistent with studies showing that donor number and cells per donor are key determinants of single-cell eQTL power^49^. Nevertheless, tissue-based single-cell studies remain essential because they capture regulatory states shaped by native vascular niches, multicellular composition and intercellular communication. Tissue-resolved QTL and perturbation studies have shown that disease-relevant genetic effects and endothelial programs can depend strongly on tissue context and cellular state^30,44,50^, while native multicellular organization provides regulatory information that is not reproduced in monoculture^51^. Our model should therefore be viewed as complementary rather than substitutive. Serial passage recapitulates a tractable form of endothelial replicative senescence, but does not fully reproduce chronological vascular aging, which is associated with endothelial senescence and vascular dysfunction^52^. Integrating donor-derived endothelial QTL maps with tissue-resolved multiomic and spatial resources will be important for assessing the generalizability of the identified regulatory effects.

Existing variant-CRE-gene prioritization frameworks are complementary but largely inferential. ABC^3^ integrates enhancer activity with promoter contact; Cicero^38^ and RobustLink^36^ infer links from co-accessibility and cross-cell covariance; supervised scE2G^39^ predicts links from single-cell features and perturbation-trained models; PHM^37^ and CIT^35^ use genetic instruments to assess putative directionality or mediation; cS2G^4^ combines multiple SNP-to-gene priors; colocalization tests^53^ shared association signals; and Hi-C and Micro-C measure physical proximity. However, physical contact does not establish regulatory function, co-accessibility may reflect shared cellular state or co-regulation, and colocalization or mediation does not by itself prove a complete variant-CRE-gene causal chain^13^. Our covariance-aware MMR occupies a distinct complementary niche by testing genotype-associated displacement of paired chromatin-expression phenotypes in the same donor, without requiring independent significance in either modality. caeQTLs should therefore be interpreted as genetically anchored joint associations rather than genome-wide mediation. Nevertheless, the significant caeQTLs reported here included not only variants with strong or moderate effects on both chromatin accessibility and gene expression, but also those with significant effects on only one trait, or even on neither trait when assessed individually, and such associations should be interpreted with caution when drawing functional conclusions. The validity of this strategy is supported by convergent evidence, including 54.28% overlapped at least one orthogonal annotation, supported sets showed greater cS2G overlap, and senescent evidence-supported caeQTLs retained CAD heritability enrichment after conditioning on caQTLs and eQTLs. Thus, caeQTLs complement existing maps by prioritizing context-dependent CRE-gene links, while residual false positives require perturbation-based validation.

Our findings extend context-dependent regulatory genetics into cellular senescence. Prior response- and trajectory-resolved QTL studies have shown that disease-associated effects can emerge or change during immune activation, differentiation and environmental perturbation^20,54,55^. In our system, senescence-associated caeQTLs provide complementary annotation of CVD loci beyond proliferating-state and single-modality maps, suggesting that part of the missing regulation may reflect the underrepresentation of disease-relevant senescent contexts rather than an absence of inherited regulatory effects^15,43^. This interpretation is necessarily incomplete because endothelial senescence represents only one vascular context, whereas CAD genetic risk is distributed across endothelial, smooth-muscle, and immune-cell states, including disease-associated SMC trajectories and macrophage-related regulatory programs^44,56,57^. Moreover, caeQTLs alone cannot resolve all downstream consequences. Histone-modification QTLs, promoter-contact QTLs, and other molecular layers can annotate additional noncoding GWAS signals and, at selected loci, support mediation, yet substantial regulatory gaps remain^10,14,58,59^. Thus, senescence-resolved caeQTLs complement existing maps by prioritizing candidate causal variants and effector genes at CVD loci.

## Methods

### Study design, sample collection and ethical approval

Human umbilical cords were collected from healthy donors after written informed consent and in accordance with protocols approved by the institutional review board of Tianjin Central Hospital of Gynecology and Obstetrics, China (approval no. 2022KY071). Primary human umbilical vein endothelial cells (HUVECs) were isolated from umbilical cord veins and expanded^60^ to establish a donor-derived cell bank of 100 genotyped, unrelated donors. This donor-scale, cell-type-resolved design was chosen to maximize statistical power for donor-level QTL mapping while minimizing the sampling biases (variable endothelial abundance, incomplete cell recovery and inter-individual differences in yield) that affect tissue-based single-cell QTL studies.

### HUVEC culture and replicative senescence model

HUVECs were cultured in Medium 199 (M199) basic medium (Gibco, C11150500BT) supplemented with 10% fetal bovine serum (FBS), 25 mM HEPES (Sigma-Aldrich, H3375), 94 µg/mL heparin sodium salt (MCE, HY-17567A), 4.17 ng/mL recombinant human β-endothelial cell growth factor (β-ECGF; Sigma-Aldrich, E1388-25UG), and 3 µg/mL thymidine (Sigma-Aldrich, T1895). Cells were maintained in culture flasks coated with 0.0029% (w/v) collagen type I from rat tail (Merck, 08-115) at 37°C in a humidified atmosphere containing 5% CO_2._ Cells at early passage (passage 1, P1) retain high proliferative capacity and normal physiological function and were defined as the proliferating state. Replicative senescence was induced by serial passaging: cells were split 1:3 during early passages and 1:2 at later passages as proliferation declined. Because the passage number required to reach replicative senescence differs among donors, senescence was instead defined functionally. Between passages 20 and 30, senescence was assessed by senescence-associated β-galactosidase (SA-β-gal) staining (Beyotime, C0602); cultures were classified as senescent when the proportion of SA-β-gal-positive cells exceeded 80%. Senescent cells also showed the expected reduction in proliferative capacity and characteristic enlarged, flattened morphology.

### Single-cell multiome (ATAC + gene expression) library generation

Matched transcriptome and chromatin accessibility were captured from the same cells using the 10x Genomics Chromium Single Cell Multiome ATAC + Gene Expression assay (10x Genomics, CG000338) according to the manufacturer’s protocol. Briefly, nuclei were isolated from each state, tagmented with Tn5 transposase, and subjected to gel-bead-in-emulsion (GEM) generation to barcode both RNA (via the gene-expression portion) and tagmented DNA (via the ATAC portion) from the same single cells. Proliferating and senescent HUVECs from all 100 donors were pooled and processed as two multiplexed libraries per state (four libraries in total). Because QTL detection power depends critically on donor number, this pooled, donor-scale design maximized the number of genotyped individuals while controlling cost. Libraries were sequenced on an Illumina NovaSeq X Plus platform with paired-end reads.

### Genotyping, quality control and imputation

Genomic DNA was extracted from each donor and genotyped on the Illumina HumanCoreExome-12 BeadChip; raw calls were generated in GenomeStudio (v2.0.5), yielding 699,533 raw variants. Genotypes were converted to PLINK binary format and filtered with PLINK (v1.9): variants with call rate <97% were removed, and individuals with a pairwise kinship coefficient >0.1875 were excluded as related, leaving 541,086 variants in the unrelated donor set. Genotypes were phased with Eagle (v2.4.1) and imputed with IMPUTE2 (v2.3.2) against the 1000 Genomes Project Phase 3 reference panel, yielding 71,568,557 imputed variants. Imputed genotypes were restricted to autosomal bi-allelic variants with minor allele frequency (MAF) >0.01 and Hardy-Weinberg equilibrium *p* >1×10^-6^, leaving 7,059,764 high-quality variants used in all genetic analyses.

### Single-cell RNA and ATAC data processing

Sequencing data were processed with Cell Ranger ARC (v2.0.2; cellranger-arc count) using the GRCh38 reference (refdata-cellranger-arc-GRCh38-2020-A-2.0.0) with default parameters, producing per-cell gene-expression and chromatin-accessibility matrices. For the RNA modality, downstream analysis was performed with Seurat (v4.4.0). Low-quality cells were filtered by the number of detected genes (2,000 ≤ nFeature_RNA ≤ 7,500) and mitochondrial read fraction (percent.mt ≤20%). Counts were normalized and variance-stabilized with SCTransform, followed by principal-component analysis, graph-based clustering and UMAP embedding. For the ATAC modality, analysis used Signac (v1.14.0). Cells were filtered on the number of fragments in ATAC peak regions (atac_peak_region_fragments), the fraction of reads in peaks (pct_reads_in_peaks), the low-mapping-quality ratio (lowmapq_ratio), nucleosome signal (nucleosome_signal) and transcription-start-site enrichment score (TSS.enrichment); data were normalized with RunTFIDF, followed by singular-value decomposition (latent semantic indexing), graph-based clustering and UMAP embedding. Because the cultures consisted of a single endothelial cell type, sparse outlier clusters were removed from both modalities as putative abnormal cells or technical noise. The two multiplexed libraries were integrated per modality with Harmony (v1.2.0) to correct batch effects while preserving donor-level signal.

### Demultiplexing of pooled cells

To assign pooled single cells to their donors of origin, we merged the RNA and ATAC BAM files to combine genetic signal across the two modalities. Vireo (v0.5.8)^34^ was run with the imputed, quality-controlled donor genotypes and the IGSR reference variant set, probabilistically assigning each cell to a donor based on the match between its allele-specific read counts and the reference genotypes. Cells classified as ’unassigned’ or ’doublet’ were discarded. Only cells confidently assigned to a known donor were retained for downstream analysis.

### Pseudo-bulk aggregation

To preserve the donor as the independent unit of genetic association, cells were aggregated within each donor and state. For the RNA modality, mean expression per gene per donor was computed from SCTransform-normalized values, followed by quantile normalization and inverse normal transformation, yielding a donor x gene matrix. For the ATAC modality, peaks were called across cells with MACS2 (v2.2.9.1); the pooled BAM was split into per-donor BAM files, reads within peaks were counted per donor with featureCounts (v2.0.6), and peaks with fewer than five total reads were removed, yielding a donor x peak count matrix.

### *cis*-eQTL mapping

*cis*-eQTLs were mapped within 1 Mb of each gene transcription start site (TSS) using a linear model implemented in BootstrapQTL (based on Matrix eQTL v1.0.5)^61^. For each gene, donor-level normalized expression E was regressed on the additive genotype dosage X_G_ and covariates:

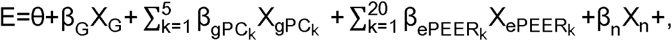

where the covariates were the top five genotype principal components (PCs), the top 20 PEER factors of the expression matrix, the reciprocal of the number of cells detected per donor (X_n_) and donor sex. PEER factors were inferred from the normalized expression matrix to capture unmeasured technical and biological confounders. Significance was assessed by local false discovery rate (LFDR), computed from the empirical null of the association statistics, with LFDR ≤0.05 considered significant. Mapping was performed separately for proliferating and senescent cells.

### *cis*-caQTL mapping

*cis*-caQTLs were mapped within 1 Mb of each peak using RASQUAL (v1.1)^62^, which models both total read count and allelic imbalance at the peak level. Covariates included the top five genotype PCs, peak GC content, the reciprocal of the number of cells detected per donor and donor sex. Empirical FDR was estimated from five permutations, and significant caQTLs were defined at FDR ≤0.05. Mapping was performed separately for proliferating and senescent cells.

### Covariance-aware joint (caeQTL) mapping

We modeled chromatin accessibility and gene expression as a paired, donor-level bivariate phenotype and tested genotype-associated displacement using multivariate multiple regression (MMR). For each candidate variant-peak-gene triplet (variant within 1 Mb of the gene TSS and within the peak; a peak-gene pair was retained only when the variant bridged them), the paired phenotypes were the donor-level normalized accessibility (A) and expression (E), and the design matrix included the covariates above (genotype PCs, expression PEER factors, peak GC content, reciprocal cell count and sex) and the additive genotype dosage G:

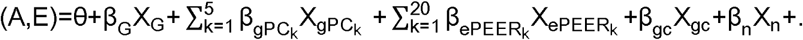

The joint null hypothesis H_0_: β_A_ (accessibility) = β_E_ (expression) = 0 was evaluated by an omnibus test comparing the full model (with G) against a reduced model (without G) using Pillai’s trace, =tr[(+)^-1^]=Σ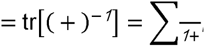, where H and E are the hypothesis and residual sum-of-squares-and-cross-products matrices and are the eigenvalues of E^-1^H. For a one-degree-of-freedom genotype term and two response variables, *V* was converted to an approximate F statistic, 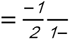, on 2 and -1 degrees of freedom (= residual degrees of freedom of the full model), yielding the joint *p*-value. Because the genotype term has a single degree of freedom, Pillai’s trace, Wilks’ lambda, Hotelling-Lawley trace and Roy’s largest root produce identical F and *p*; we report Pillai’s trace *V* as a non-directional measure of joint effect magnitude (0 ≤ V ≤ 1). The nested-model comparison was implemented in R using lm(cbind(A, E) ∼ covariates + G) and anova(fit0, fit1, test = “Pillai”) so that the genotype effect is assessed as a partial test after adjustment for all covariates.

*P*-values were adjusted across all tested triplets within a state by the Benjamini-Hochberg (BH) procedure, and associations with adjusted *p* ≤0.05 were termed chromatin accessibility-expression QTLs (caeQTLs). This triplet-level BH controls the variant-peak-gene FDR, which is distinct from variant-, feature- or locus-level FDR. For each significant caeQTL we additionally report the component-specific statistics (β_A_, *p*_A_; β_E_, *p*_E_).

We note explicitly that a significant joint association indicates that the genotype affects at least one of the two molecular phenotypes, and does not by itself establish that accessibility mediates the effect of genotype on expression; caeQTLs therefore denote genetically anchored joint associations rather than genome-wide mediation. To distinguish dual-modality from single-modality joint effects, the component statistics (β_A_/β_E_ and their *p*-values) should be examined alongside the omnibus *V*.

### Functional annotation of QTL variants

QTL variants were annotated against ENCODE V4 candidate cis-regulatory elements (cCREs)^63^ and classified as promoter-like signatures (PLS), proximal enhancer-like signatures (pELS), distal enhancer-like signatures (dELS), CTCF-bound (CA-CTCF), H3K4me3-modified (CA-H3K4me3), TF-bound (CA-TF), chromatin-accessible-only (CA) or TF-only regions, or as non-cCRE. Variants were additionally annotated by gene-region structure (intron, 5′ UTR, 3′ UTR). Enrichment of significant QTL variants relative to non-significant variants within each annotation was computed as the ratio of the fraction of QTL variants to the fraction of non-QTL variants overlapping that annotation.

### QTL state-specificity classification

QTLs significant (FDR <0.05) in both states were classified as state-shared; QTLs significant in only one state were classified as state-inclined. This FDR-based classification was cross-validated with Cochran’s Q heterogeneity test, in which QTLs with Cochran’s Q *p* >0.05 and I² <25% were considered state-shared and the remainder state-inclined.

### Colocalization analysis

Cross-state colocalization of eQTLs and caQTLs was assessed with coloc (v5.2.3)^64^, using all variants per feature, with significant colocalization defined as PP4 ≥0.75 and PP4/PP3 ≥3; 95% credible sets were derived where applicable. Multi-phenotype/multi-signal colocalization (eQTL-caQTL, and later caQTL/eQTL with GWAS) was assessed with moloc (v0.1.0)^65^ using default parameters, with posterior probability ≥0.8 taken as evidence of colocalization. To reduce computational burden, eQTL-caQTL colocalization was restricted to gene-peak pairs supported by functional evidence. Colocalization results were visualized with locuscomparer (v1.0.0)^66^.

### CRE-gene linking and causal evidence

Peak-gene regulatory relationships were assembled from multiple orthogonal methods: RobustLink (|correlation| >0.25)^36^, the Pairwise Hierarchical Model (PHM; posterior probability of causality ≥0.5)^37^, the Causal Inference Test (CIT v2.3.2; TassocGgvnL *p* <0.05)^35^, Cicero (|co-accessibility score| >0.1)^38^, the Activity-by-Contact model (ABC; default score threshold 0.013)^3^, and scE2G (quantile-normalized E2G score threshold 0.164)^39^. These were complemented by chromatin-loop evidence from Hi-C data for HUVECs obtained from the 3D Genome Browser^67^ and in-house HUVEC Micro-C data. Each source provides an independent, complementary form of support (physical contact, co-accessibility/covariance, or genetic causality), and a caeQTL feature was considered evidence-supported if at least one source endorsed its peak-gene link.

### Epigenomic state enrichment

Enrichment of QTL variants across the 25 chromatin states of the Roadmap HUVEC line (E122) was evaluated by intersecting significant QTL variants (positive set) and non-significant variants (negative set) with each state annotation using bedtools (v2.30.0), and computing the odds ratio of overlap between the two sets. The 25 states span promoter, enhancer, transcriptional elongation and heterochromatic categories.

### Fine-mapping

Significant eQTL and caQTL loci were fine-mapped with CAVIAR^40^, allowing up to three causal variants per locus (maximum causal variants = 3); variants with causal posterior probability >0.8 were retained as high-confidence candidates. For caeQTLs, candidate variants were prioritized by the same Bayesian framework applied to the joint signal.

### Transcription-factor footprinting

TF binding dynamics were assessed from single-cell ATAC data with TOBIAS (v0.14.0)^68^. After merging the 215,778 peaks called across both states, the ATACorrect module was used to correct Tn5 sequence bias, ScoreBigwig to compute per-position footprint scores, and BINDetect to predict TF occupancy. Motif collections were obtained from JASPAR 2024 (2,346 motifs)^41^ and HOCOMOCO v13 (1,611 motifs)^42^. Differential binding between states was summarized by a change score (difference in binding score) and a background-model *p*-value; candidate TFs were flagged when -log_10_(*p*) exceeded the 95th percentile across all TFs and/or the change score fell below the 5th or above the 95th percentile.

### Gene-set enrichment analysis

KEGG pathway enrichment was performed with clusterProfiler (v4.15.2) using a hypergeometric test, with BH-corrected *p* <0.05 as the significance threshold.

### Stratified LD score regression (S-LDSC)

Heritability enrichment of caeQTL, caQTL, and eQTL variant sets for cardiovascular-disease (CVD) traits was assessed with S-LDSC (ldsc, v1.0.1)^69^, using the 1000 Genomes Phase 3 East Asian (EAS) LD reference panel to match the LD structure of the target population and CVD GWAS summary statistics. Annotation sets were constructed from the three QTL variant classes without additional gene annotation; enrichment scores and their significance were computed per annotation. Conditioned analyses additionally included caQTL and eQTL annotations in the baseline model.

### π_1_ statistic

Replication of HUVEC QTLs against GTEx arterial eQTLs was quantified with the π_1_ statistic. For each tissue, the *p*-value distribution of HUVEC variant-gene pairs in GTEx was used to estimate the null fraction π_0_ with qvalue (v2.1.1), and π_1_ = 1 - π_0_ was computed as the fraction of non-null (true) associations; larger π_1_ indicates greater replication.

### GWAS overlap and SNP-to-gene prioritization

Disease-relevant variants were obtained from the CAUSALdb2 database^2^ across eight disease systems. cS2G scores^4^ were used to assess SNP-to-gene support of QTL variants overlapping GWAS signals, and CVD GWAS credible sets were used to benchmark QTL-prioritized variants. Cross-state sharing with GTEx^70^ was quantified with

Storey’s π_1_ as described above.

### Chromatin accessibility and histone modification profiling

The chromatin accessibility and histone modification (H3K27ac, H3K4me1, and H3K4me3) profiling data for proliferating, intermediate, and senescent HUVECs were obtained from the Gene Expression Omnibus (GEO)^71^. Read quality was assessed with FastQC (v0.12.0) and reads were trimmed with Trim Galore (v0.6.10), aligned to hg38 with Bowtie2 (v2.4.3), deduplicated with Picard MarkDuplicates (v3.4.0), and peaks were called with MACS2 (v2.2.9.1).

### Allele-specific ATAC-PCR

To quantify allele-specific chromatin accessibility at rs2019090, heterozygous cells were subjected to the ATAC transposition reaction and library preparation described above. The rs2019090-containing region was then amplified from the ATAC library with primers flanking the SNP (forward: 5’-TACCATTACTGGTGGGTCAG-3’; reverse: 5’-GGTCACTATCTGGTGGTTTAT-3’), and the amplicons were Sanger-sequenced. Allelic imbalance was quantified from the relative A:T peak heights at rs2019090; in heterozygous cells, deviation from a 1:1 A:T ratio reflects allele-specific chromatin accessibility, and the imbalance was compared between proliferating and senescent cells.

### Dual-luciferase reporter assay

A ∼500-bp fragment flanking rs2019090 was amplified from genomic DNA by overlap-extension PCR to introduce the A or T allele, and cloned into the pGL3-SV40 reporter vector upstream of the SV40 promoter. Outer primers were rs2019090-F (5’-ATTTCTCTATCGATAGGTACCTATAGAAAAGGATTTCCAGCAA-3’) and rs2019090-R (5’-ACTAATTGAGATGCACTCGAGGCTGTCCAGTCGCAAACACA-3’). Allele-specific inner primers were rs2019090-A-F (5’-GCCTAAATTTATTTTCTTAAA-3’) and rs2019090-A-R (5’-TTTAAGAAAATAAATTTAGGC-3’) for the A allele, and rs2019090-T-F (5’-GCCTAAATTTTTTTTCTTAAA-3’) and rs2019090-T-R (5’-TTTAAGAAAATAAATTTAGGC-3’) for the T allele. Sanger-verified plasmids were co-transfected with pRL-TK Renilla luciferase into 293T cells in 24-well plates using LipoFiterTM Liposomal Transfection Reagent (HANBIO, HB-LF-1000); luciferase activities were measured 48h later with the Dual-Luciferase Reporter Assay System, and firefly luciferase was normalized to Renilla. Relative luminescence reflected the enhancer activity of each allele. Parallel experiments were conducted in HUVECs following the identical protocol.

### Electrophoretic mobility shift assay (EMSA)

Biotin-labelled double-stranded probes encompassing rs2019090 were annealed by heating to 95°C for 15 min followed by gradual cooling to room temperature. Probe sequences were: BIO-rs2019090-A-F (5’-CTTTGGTCTAGCCTAAATTTATTTTCTTAAAGTTTAAGCTC-3’), BIO-rs2019090-A-R (5’-GAGCTTAAACTTTAAGAAAATAAATTTAGGCTAGACCAAAG-3’), BIO-rs2019090-T-F (5’-CTTTGGTCTAGCCTAAATTTTTTTTCTTAAAGTTTAAGCTC-3’) and BIO-rs2019090-T-R (5’-GAGCTTAAACTTTAAGAAAAAAAATTTAGGCTAGACCAAAG-3’). Nuclear extracts were prepared from HUVECs using a nuclear-protein extraction kit. Labelled probes (20 nM) were incubated with nuclear extracts in binding buffer at 4°C for 1 h. Protein-DNA complexes were resolved on a non-denaturing polyacrylamide gel in 0.5 x TBE at 100 V for 1.5 h, transferred to a nylon membrane at 380 mA for 30 min, UV cross-linked, and detected with HRP-conjugated streptavidin and chemiluminescence. Allele-specific binding was assessed by comparing the shifted-band intensities of the A and T probes.

### Single-nucleotide editing

The endogenous A allele of rs2019090 was edited to T in immortalized HUVECs by CRISPR-Cas9-mediated homology-directed repair (HDR). A sgRNA targeting the locus was designed with CRISPOR (sgRNA oligos: 5’-CACCGCATTACACAGACCTCAGTC-3’ and 5’-AAACGACTGAGGTCTGTGTAATGC-3’) and cloned into pX459. The sgRNA plasmid and a single-stranded oligodeoxynucleotide (ssODN) repair template carrying the T allele were co-transfected at a 1:2 plasmid:template ratio. After 6h the medium was replaced, and after 24 h puromycin (Solarbio, P8230) (1.5 µg/mL) was added for selection until control cells died. Single-cell clones were isolated by flow-cytometric sorting into 96-well plates, expanded, and genotyped by PCR amplification of the edited locus (genotyping primers: 5’-TACCATTACTGGTGGGTCAG-3’ and 5’-GGTCACTATCTGGTGGTTTAT-3’) followed by Sanger sequencing. Clones confirmed as homozygous-edited were used for downstream functional assays.

### DNA pull-down and mass spectrometry

Biotin-labelled A- and T-allele probes (sequences as in the EMSA section) were annealed and immobilized on streptavidin magnetic beads after blocking. Nuclear extracts were pre-cleared to reduce non-specific binding, then incubated with the probe-bead complexes at 4°C with rotation. Beads were washed with a graded series of wash buffers, and bound proteins were eluted, resolved by SDS-PAGE, stained with a mass-spectrometry-compatible silver stain, and the A- and T-specific gel regions were excised. After in-gel reduction (DTT), alkylation (IAA) and trypsin digestion, peptides were desalted on a C18 microcolumn and identified by LC-MS/MS. Allele-specific enrichment was determined from spectral counts and sequence coverage, comparing proteins captured by the A and T probes.

### RT-qPCR

Total RNA was extracted with TRIzol (phenol-chloroform), and RNA concentration and purity (A_260_/A_280_) were assessed by NanoDrop. One microgram of total RNA was reverse-transcribed to cDNA, and qPCR was performed with SYBR Green on a real-time PCR system (95°C initial denaturation followed by 40 cycles of denaturation, annealing and extension). GAPDH served as the internal control, and relative expression was calculated by the 2^-ΔΔCt^ method. Primers were: qGAPDH-F (5’-GGAAGGTGAAGGTCGGAGTCA-3’), qGAPDH-R (5’-GTCATTGATGGCAACAATATCCACT-3’), qHMGA1-F (5’-GCTGGTAGGGAGTCAGAAGGA-3’), qHMGA1-R (5’-TGGTGGTTTTCCGGGTCTTG-3’), qPDGFD-F (5’-CAACCTCAGGCGAGATGAGA-3’) and qPDGFD-R (5’-GGTTCCTGGGGTAGCTGTTC-3’).

### Allele-specific MNase-based 4C

For allele-specific MNase-based 4C analysis, approximately 1 x 10^7^ cells heterozygous for the rs2019090 A/T variant were collected. Chromatin was subjected to dual cross-linking with 1% formaldehyde (ThermoFisher, 28906) for 10 min followed by 3 mM disuccinimidyl glutarate (DSG, ThermoFisher, 20593) for 45 min. Nuclei were subsequently isolated using a lysis buffer containing NP-40. Chromatin was digested with micrococcal nuclease (MNase, Worthington Biochem, LS004798), and an MNase titration experiment was performed using the same batch of samples to determine the optimal digestion condition, targeting a digestion profile of approximately 80-90% mononucleosomes and 10-20% dinucleosomes. Following MNase digestion, DNA ends were end-repaired using T4 polynucleotide kinase (T4 PNK, New England BioLabs, M0201) and Klenow fragment (New England BioLabs, M0210). The repaired DNA ends were subsequently labeled with biotin-dATP (Jena Bioscience, NU-835-BIO14), biotin-dCTP (Jena Bioscience, NU-809-BIOX), dTTP (ThermoFisher, 10219012), and dGTP (ThermoFisher, 10218014), followed by proximity ligation using T4 DNA ligase (New England BioLabs, M0202). Biotin nucleotides at unligated DNA ends were removed using exonuclease III (New England BioLabs, M0206). The samples were then treated with proteinase K (Vazyme, DE102-01) and SDS (Sigma-Aldrich, L3771) to reverse the cross-links, followed by DNA extraction. The resulting DNA was size-selected by electrophoresis on a 1.5% agarose gel, and DNA fragments larger than 300 bp were recovered. Libraries were subsequently prepared using a 4C-based PCR approach with 2 x Phanta Max Master Mix (Vazyme,P525) and a viewpoint-specific primer targeting the rs2019090-containing region. Specifically, a US primer (5’-AATGATACGGCGACCACCGAGATCTACACTCTTTCCCTACACGACGCTCTTCCGATCTAACCACGGGTTTGCAACATT-3’) and a universal primer (5’-CAAGCAGAAGACGGCATACGA-3’) were used for PCR amplification. The resulting libraries were subjected to sequencing for allele-specific chromatin interaction analysis. Reads were trimmed with Cutadapt (v5.1), aligned uniquely to hg38 with Bowtie2 (v2.4.3), deduplicated, and 4C interaction profiles were quantified with custom scripts.

### Motif scanning

The 25-bp sequence flanking rs2019090 (5’-TAGCCTAAATTTA(T)TTTTCTTAAAGT-3’, where A is the risk allele and T in parentheses the protective allele) was scanned with FIMO (v5.5.9)^72^ against the JASPAR 2024 motif database^41^; matches with *p* <0.01 and *q* <0.05 were retained for each allele.

### Western blotting

Cells were lysed in RIPA buffer with protease inhibitors, and protein concentrations were determined by a protein quantification kit. Equal amounts of protein were resolved by SDS-PAGE (80V stacking, 120V resolving) and wet-transferred to a PVDF membrane (activated in methanol). Membranes were blocked, probed with primary antibodies against HMGA1, PDGFD and GAPDH (4°C, overnight), washed, incubated with HRP-conjugated secondary antibody, and developed with ECL. Band intensities were normalized to the loading control (GAPDH) for quantification.

### CUT&Tag assay

Approximately 1 x 10^5^ HUVECs were collected and immobilized on VAHTS Concanavalin A-coated Magnetic Beads Pro (Vazyme, N515). The bead-bound cells were incubated with an anti-HMGA1 antibody (abcam, ab129153) at 4°C overnight, followed by incubation with the corresponding Rabbit IgG (H&L) Antibody Pre-Adsorbed (Rockland, 611-201-122) at room temperature for 1 h. The cells were subsequently incubated with pA/G-Tn5(Vazyme, S604) transposase at room temperature for 1 h. Tagmentation was initiated by adding MgCl_2_-containing Tagmentation Buffer and incubating the samples at 37°C for 1 h, allowing targeted chromatin fragmentation and adapter integration. The reaction was then terminated, and the samples were treated with SDS, and proteinase K to facilitate protein digestion and DNA release. DNA was subsequently extracted using phenol:chloroform:isoamyl alcohol (25:24:1) (JSENB, JS0700). The purified tagmented DNA was amplified using TruePrep® Amplify Enzyme (Vazyme, TD601) to generate sequencing libraries, which were purified and subjected to paired-end 150-bp sequencing on an Illumina platform. Read quality was assessed with FastQC (v0.12.0) and reads were trimmed with Trim Galore (v0.6.10), aligned to hg38 with Bowtie2 (v2.4.3), deduplicated, and signal was normalized to counts per million (CPM) with deepTools (v3.5.6).

## Acknowledgements

The work was supported by the following grants: the National Natural Science Foundation of China (82422038 to D.D.H., 32470719 to X.F.Y. and 32470671 to M.J.L.), and the Natural Science Foundation of Tianjin (24JCZDJC00480 to M.J.L.).

## Author contributions

M.J.L. and X.F.Y. conceived and designed the study, supervised the project, and wrote the manuscript. X.F.Y., W.D. and Y.Z. performed the data analysis, developed the computational framework, and wrote the manuscript. X.Q.W., K.L.L., L.Z., F.C., Q.T.J., H.L., Y.C. and X.Q.G. collected the clinical samples, performed the functional experiments, and validated the key findings *in vitro*. X.F.Y., J.H.W., W.D., Y.Z. and D.D.H. contributed to data processing, statistical analysis, and interpretation of results. P.C.S. and D.D.H. contributed scientific expertise and statistical guidance, and critically reviewed the manuscript. All authors read and approved the final manuscript.

## Data availability

The raw sequence data reported in this paper have been deposited in the Genome Sequence Archive in National Genomics Data Center, China National Center for Bioinformation/Beijing Institute of Genomics, Chinese Academy of Sciences (GSA-Human) that are publicly accessible at https://ngdc.cncb.ac.cn/gsa-human.

## Supplementary Figure Legends

**Fig. S1 | Single-cell eQTL and caQTL detection during endothelial senescence. A**, UMAP of scRNA-seq (top) and scATAC-seq (bottom) data in proliferating (left) and senescent (right) cells. **B**, QQ plots of eQTLs (top) and caQTLs (bottom) in both states. **C**, Overlap of significant QTLs between states at the feature (eGene/caPeak), variant, and association levels (top, eQTL; bottom, caQTL). **D**, Comparison of eQTL effect sizes with GTEx Artery-Coronary (top) and Artery-Tibial (bottom) eQTLs. **E**, Replication of HUVEC eQTLs in GTEx Artery-Aorta/Coronary/Tibial tissues at variant, eGene, and variant-eGene levels. **F**, Reverse comparison: the proportion of GTEx eQTLs detected in HUVECs. **G**, Proportion of GTEx eQTLs among proliferating-inclined, senescent-inclined, and shared HUVEC eQTLs. **H**, Genome-wide differential accessibility between proliferating and senescent cells around ATAC peaks.

**Fig. S2 | Cross-validation of state-specific QTL classification. A**, Agreement between FDR-threshold and Cochran’s Q classifications of caQTL state specificity. **B**, As in A, for eQTLs. **C**, Effect-size comparison between concordant and discordant caQTL classifications (Cochran’s Q). **D**, Comparison of caQTL effect sizes by classification agreement within FDR-defined groups. **E**, As in **C**, for eQTLs. **F**, As in **D**, for eQTLs.

**Fig. S3 | Colocalization analysis of HUVEC QTLs. A**, Representative cross-state eQTL colocalization: a shared but non-colocalized example (*ZFP30*) and a proliferating-inclined colocalized example (*RGS12*, rs13104157). **B**, Representative caQTL colocalization: a shared but non-colocalized peak (chr15:85296348-85297292) and a senescent-inclined colocalized peak (chr1:203152220-203154745, rs149200581). **C**, Distance of significant eQTL and caQTL variants from TSS. **D**, Overlap of significant eQTL and caQTL variants in proliferating (left) and senescent (right) cells. (Top) Venn diagrams show the numbers of variants significant only as eQTLs (eQTL-specific), only as caQTLs (caQTL-specific), or as both (shared). (Bottom) For each cell state, bars indicate the percentage of variants in each class with CIT evidence.

**Fig. S4 | Fine-mapping of caQTLs and eQTLs. A**, Summary of fine-mapping results: the distribution of candidate causal variants per caPeak (left) and eGene (right) in proliferating and senescent cells. **B**, Multi-dimensional comparison of fine-mapped molecular-QTL variants in proliferating (top) and senescent (bottom) cells, showing overlap between fine-mapped variants, significant QTL variants (caQTL and eQTL), and variants with CIT support. **C**, Epigenomic-state enrichment of fine-mapped eQTL and caQTL variants across the 25 Roadmap HUVEC (E122) chromatin states, recapitulating the enrichment pattern of the full QTL maps.

**Fig. S5 | Transcription-factor footprinting across proliferating and senescent cells. A**, JASPAR 2024 footprinting overview: differential binding score (x-axis) versus statistical significance (y-axis, −log_10_(*p*) from the background model) for each TF. **B**, As in A, using HOCOMOCO v13 motifs. **C**, Differential TF binding between proliferating and senescent cells from HOCOMOCO v13 motifs; TFs are flagged when −log_10_(*p*) exceeds the 95th percentile and/or the differential binding score falls outside the 5th/95th percentiles (\*\*\**p* <1e-200, \*\**p* <1e-150, \**p* <1e-100). Significance and the 5th/95th-percentile change thresholds are defined in Methods.

**Fig. S6 | caeQTL detection from single-cell multiome data. A**, QQ plots of caeQTLs in proliferating (top) and senescent (bottom) states. **B**, Manhattan plots of caeQTLs. **C**, Overlap of significant caeQTLs between states at feature (peak-gene pair), variant, and association levels. **D**, Composition of caeQTLs by the combination of accessibility and expression effects; many significant joint associations lack individually significant effects in either modality.

**Fig. S7 | Systematic comparison of caeQTLs with caQTLs and eQTLs. A**, Number of input (left) and significant output (right) tests at variant, feature, and association levels. **B**, Distribution of variants per feature among inputs (left) and significant outputs (right). **C**, Overlap of significant caeQTLs with eQTLs (top) and caQTLs (bottom) at variant (left) and association (right) levels.

**Fig. S8 | caeQTL gains power from the joint information of two phenotypes. A**, Proportion of caeQTLs that are also significant caQTLs or eQTLs. **B**, Composition of caeQTLs by corresponding QTL class. **C**, Overlap of significant results among caeQTL, caQTL, and eQTL. **D**, *P*-value distributions of the three QTL classes. **E**, Power curves of the three QTL classes stratified by effect magnitude.

**Fig. S9 | Functional annotation of caeQTLs. A**, Distance of caeQTL and caQTL variants from TSS. **B**, KEGG enrichment of genes preferentially highlighted by joint rather than expression-only evidence. **C**, KEGG enrichment comparison between caeQTL- and eQTL-linked genes. **D**, Comparison of effect estimates between eQTLs corresponding to significant caeQTLs and GTEx arterial eQTLs. **E**, Replication of caeQTL variant-gene pairs in GTEx eQTLs (variant-gene, variant, and eGene levels). **F**, Logistic-regression assessment of genomic and epigenomic predictors of caeQTL status.

**Fig. S10 | Colocalization and prioritization of caeQTLs. A**, Overlap between proliferating and senescent states for caeQTL-specific, shared, and colocalization-specific associations. **B**, Overview of caeQTL prioritization, showing overlap between prioritized variants, significant QTL variants (caeQTL, caQTL, and eQTL) and variants with CIT support.

**Fig. S11 | Association of caeQTLs with GWAS. A**, Difference in GWAS-variant overlap of caeQTLs versus caQTLs and eQTLs (top, proliferating; bottom, senescent). **B**, Composition of GWAS-overlapping QTL variants by caeQTL/caQTL/eQTL class across systems. **C**, cS2G support for GWAS-overlapping QTL variants across QTL classes and state categories. **D-G**, Comparison of CVD GWAS-overlapping variants across caeQTL, caQTL, and eQTL and across states, shown as state-stratified Venn diagrams of fine-mapped (**D**) or all significant (**E**) variants, and as overall UpSet plots of all significant (**F**) or fine-mapped (**G**) variants combining both states and the three QTL types. **H**, Replication of caeQTL, caQTL, and eQTL variants across eight disease systems in proliferating (top) and senescent (bottom) cells. **I**, Ranked replication of caeQTL (left), caQTL(middle), and eQTL (right) variants across eight disease systems, grouped by state specificity.

**Fig. S12 | Comparison of caeQTL, caQTL and eQTL colocalization with CVD GWAS. A**, Overall comparison (*, significant in caeQTL analysis; PPA, posterior probability of association). **B**, Representative examples (green, colocalization-only; purple, caeQTL-only).

**Fig. S13 | QTL signals for CAD-relevant variant-to-gene-to-program (V2G2P) genes. A**, *PLPP3* single-cell signals across genotypes (left, *PLPP3*-associated peak accessibility; right, *PLPP3* expression). **B**, Bulk epigenetic signal of the *PLPP3*-associated peak across states. **C**, *PECAM1* across caeQTL, caQTL, and eQTL in proliferating and senescent cells (*, statistically significant). **D**, *COL4A2* across caeQTL, caQTL, and eQTL in proliferating and senescent cells (*, statistically significant).

**Fig. S14 | Molecular mechanism of rs2019090-mediated regulation. A**, Effect size and significance of rs2019090 in eQTL analysis (*, statistically significant). **B**, −log_10_(*p*) of rs2019090 across QTL classes. **C**, Genomic relationship among rs2019090, its peak (chr11:103797667-103798761), and *PDGFD*. **D**, ATAC signals and histone-modification (H3K27ac, H3K4me1, and H3K4me3) at the rs2019090 peak across states. **E**, Dual-luciferase reporter assay in 293FT cells showing allele-dependent enhancer activity. **F**, Migration of HUVECs across genotypes.

**Fig. S15 | HMGA1-dependent regulation of *PDGFD* by rs2019090. A**, MNase-based 4C comparing rs2019090-*PDGFD* promoter interaction strength across states. **B**, Mass-spectrometry comparison of DNA pull-down products between A and T alleles, showing allele-unique and shared bound proteins. **C**, Motif scanning of A- and T-allele flanking sequences. **D**, Correlation of *HMGA1* and *PDGFD* expression in GTEx arterial tissues (aorta, coronary).

### Supplementary Table Legends

Table S1: eQTLs in proliferating HUVECs Table S2: eQTLs in senescent HUVECs

Table S3: caQTLs in proliferating HUVECs Table S4: caQTLs in senescent HUVECs

Table S5: eQTL colocalization between proliferating and senescent HUVECs

Table S6: caQTL colocalization between proliferating and senescent HUVECs

Table S7: eQTL-caQTL colocalization in proliferating HUVECs

Table S8: eQTL-caQTL colocalization in senescent HUVECs

Table S9: caQTL fine-mapping results

Table S10: eQTL fine-mapping results

Table S11: caeQTLs in proliferating HUVECs Table S12: caeQTLs in senescent HUVECs

Table S13: Evidence support for caeQTL features

Table S14: Prioritized caeQTL results for proliferating HUVECs

Table S15: Prioritized caeQTL results for senescent HUVECs

Table S16: Significant colocalization among caQTL, eQTL and CVD GWAS

Table S17: Mass-spectrometry results for rs2019090 A- and T-allele probes

Table S18: Motif-scanning results for rs2019090 A and T alleles

